# 3D-bioprinted patient-specific organotypic bone model mimicking mineralization dysregulation in *FKBP10*-related osteogenesis imperfecta

**DOI:** 10.1101/2024.05.20.594917

**Authors:** Julia Griesbach, Anke de Leeuw, Tanja Minacci, Ben Kodiyan, Timothée Ndarugendamwo, Pei Jin Lim, Marianne Rohrbach, Marina Rubert, Matthias Rüger, Cecilia Giunta, Friederike A. Schulte, Ralph Müller

**Affiliations:** Institute for Biomechanics, ETH Zurich; Zurich, Switzerland; Connective Tissue Unit, Division of Metabolism and Children’s Research Center, University Children’s Hospital Zurich; University of Zurich, Zurich, Switzerland; Department of Pediatric Orthopaedics and Traumatology, University Children’s Hospital Zurich; Zurich, Switzerland

## Abstract

Osteogenesis imperfecta (OI) is a heterogeneous group of rare genetic diseases characterized by increased bone fragility and deformities. The pathomechanisms of OI are poorly understood, hindering the development of disease-specific therapy. Addressing the limited understanding of OI and the lack of targeted treatments remains a challenge, given its varied symptoms and large clinical spectrum. Animal models have greatly advanced the understanding of the disease; however, the heterogeneity and subtype-specific symptoms are difficult to translate to humans. *In vitro* models offer a promising tool for translational medicine, as they have the potential to yield patient-specific insights in a controlled environment using patient derived-cells. We used mechanically loaded 3D-bioprinted patient-specific organotypic bone models and time-lapsed micro-computed tomography to demonstrate dysregulation of mineralization in *FKBP10*-related OI compared to healthy controls. In contrast to healthy controls, tissue mineral density and stiffness were decoupled, such that hypermineralization observed in OI samples did not lead to increased stiffness. Additionally, we were able to replicate experimental stiffness using sample specific micro-finite element analysis. This allowed us to show mineral formation in regions of high local strain, suggesting mechanoregulation in *FKBP10*-related OI organotypic bone models is comparable to healthy controls. Regional analysis of mineralization showed increased heterogeneous mineralization, microarchitectural inhomogeneities and scaffold microporosity of OI samples compared to healthy controls. Our results suggest that the observed dysregulation of mineralization is the main driver for the altered mineral-mechanics properties observed in *FKBP10*-related organotypic bone models.

**One Sentence Summary:** Organotypic bone models demonstrate dysregulated mineralization in osteogenesis imperfecta samples compared to healthy controls.

## INTRODUCTION

Osteogenesis imperfecta (OI), commonly known as brittle bone disease, describes a spectrum of genetic skeletal disorders characterized by bone fragility and deformities. With an incidence of 1 in 10-20,000 births, OI is one of the most prevalent skeletal conditions among rare diseases (*1, 2*). The complexity and heterogeneity of OI are underlined by its classification into over 20 distinct types (*1, 3, 4*).While the most common causes for OI stem from dominant mutations in *COL1A1* and *COL1A2* genes, encoding the collagen type I α1 and α2 chains, many of the additional causative genes show autosomal as well as x-linked recessive inheritance patterns (*5*). Phenotypic variability exists within each genetic form of OI, with a spectrum of different clinical presentation observed across patients with different pathogenic variants and also among family members carrying the same variant (*6–8*). This geno- and phenotypic heterogeneity illuminates the diversity of mechanisms underlying OI pathogenesis, including impaired collagen synthesis, post-translational modifications, and compromised osteoblast function (*1, 9*).

While OI is characterized by low bone mass, bone mineralization is increased independently of the underlying specific collagen mutation and structure (*10, 11*). This is hypothesized to contribute to the ‘brittleness’ of the bones (*10, 12, 13*). Quantitative backscattered electron imaging (qBEI) analyses on bone biopsies from pediatric OI cases revealed consistent abnormal mineralization density patterns irrespective of clinical severity (*14–17*), with bisphosphonate treatment showing no additional increase in bone matrix mineral content, presumably due to inherent mineral saturation within the bone matrix (*18*). Bone mineral density (BMD) in the patients has been investigated using dual-energy X-ray absorptiometry (DXA) and high-resolution peripheral quantitative computed tomography (HR-pQCT). However, usually BMD is evaluated as areal BMD or volumetric BMD, which are average parameters influenced by porosity, cortical thickness, or trabecular volume (*12*). Most studies have found lower areal BMD in OI patients, independent of age (*19, 20*), however, volumetric BMD studies have found high variations within one patient and even the same bone ranging from lower BMD than healthy control, to similar values, or even higher BMD than healthy control (*21–23*), suggesting these standardized average parameters might not fully capture the heterogeneity of bone mineralization of OI. Additionally, a recent study found microarchitectural inhomogeneities in OI, with both sclerotic and void areas in trabecular bone, which were not always reflected in the quantitative results of the analysis (*24*), but likely contribute to poor bone quality (*25*). The exact causes for brittle bones in OI are still not fully understood, contributing to why treatment of OI remains challenging. With no curative interventions available, disease management is focused on supportive measures tailored to individual needs (*9, 26*). Given the spectrum of severity, degree of disability, and broad age range within OI populations, a multidisciplinary and personalized approach to treatment is imperative.

Animal models have been extremely valuable to study disease pathomechanisms and test therapeutic interventions (*27, 28*). However, while often informative, clinical manifestations can vary between species due to differences in bone anatomy and physiology (*29, 30*). The generation of transgenic animals and their husbandry also involve high costs, and existing animal models represent only about half of the genetic forms of human OI. Notably, *FKBP10*-related OI (*31*) is a moderately severe form of OI resulting from the faulty co/post translational processing of collagen type I, characterized by bone fragility and joint contractures (*6, 32*). *FKBP10* knock-out mice died prenatally (*33*), while other conditional models only partially recapitulated the human condition, with mild bone phenotypes (*34, 35*). To overcome the limitations of animal models, *in vitro* studies of patient-derived cells show great potential. In monolayer cell cultures, *FKBP10*-related OI fibroblasts showed a delay in collagen secretion, reduced collagen crosslinking and disorganized, branched collagen matrices (*36*). However, these 2D cultures fail to reproduce the complexity of bone tissue or capture mineralization phenotypes in OI. 3D *in vitro* bone models may preserve patient-specific attributes while enhancing complexity and functionality beyond monolayer cultures, enabling comprehensive OI research. A recently developed osteogenic organoid modeling a lethal type II OI using induced pluripotent stem cells (iPSC) demonstrated the potential of such models to gain insights into disease mechanisms (*37*). However, iPSC induction and differentiation procedures are time and labor-intensive and may introduce (epi)genetic variation compared to the original primary cells from which they were derived, posing a challenge for disease modelling (*38*). Implementing an OI model using patient-specific primary osteoblasts offers a straightforward approach, reducing study duration and minimizing the risk of off-target cell types.

In this study, our goal was to develop a patient-specific organotypic bone model for bone mineralization with *FKBP10*-related OI as proof-of-concept, here simply referred to as OI. Using primary osteoblasts derived from an OI patient and a metabolically healthy control, we employed 3D bioprinting technology to fabricate the bone constructs (Fig. 1). The constructs were physiologically loaded in a compression bioreactor and evaluated by weekly time-lapsed micro-computed tomography scans. Sample-specific micro-finite element (micro-FE) modelling allowed the assessment of mechanoregulation, the influence of the local mechanical environment on mineralization. We further defined regions of the scaffolds to investigate mineralization heterogeneity and microarchitecture in OI. Assessment of organotypic bone models using spatially correlated histological and micro-CT sections supported the detailed characterization of the dysregulated mineralization. We expect the organotypic bone model will facilitate further study of the complex and variable nature of osteogenesis imperfecta through integrated *in vitro* and *in silico* approaches, offering insights into the pathogenesis of the disease and potential therapeutic targets.

**Fig. 1.**
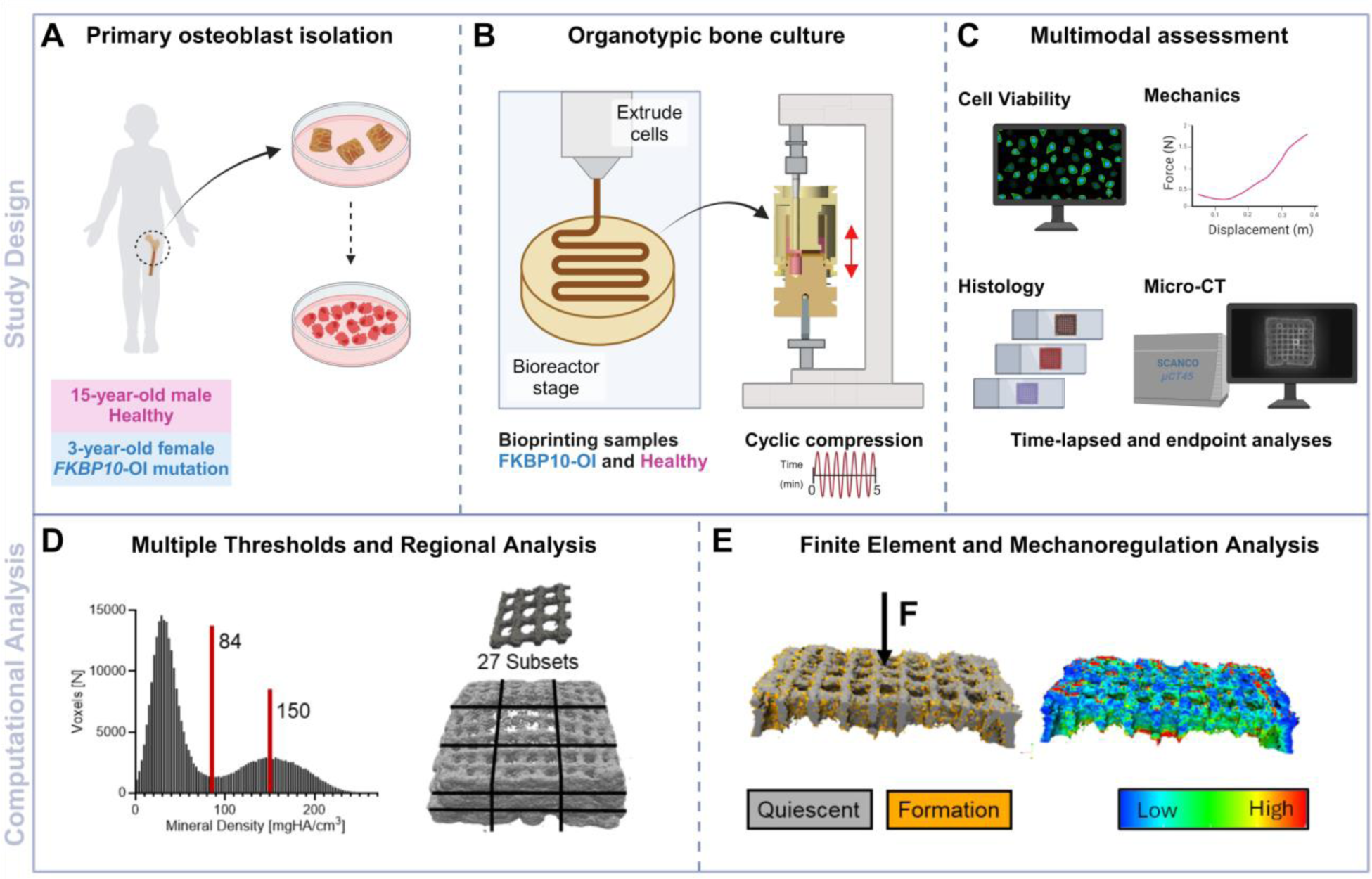
Study design and analysis. (**A**) Bone was material is collected from patient undergoing orthopedic surgery. Primary osteoblasts are isolated from donor bone, expanded to increase cell numbers, (**B**) and combined with an osteogenic bioink (0.8% alginate, 4.1% gelatin, 0.1% graphene oxide microparticles) for extrusion bioprinting. Organotypic bone models are cultured under cyclic compressive loading for 8 weeks (5 mins, 5x/week, 1% strain, 5Hz). (**C**) Bioprinted samples undergo multimodal time-lapsed and endpoint analysis, investigating cell viability, histology, and mechanical properties to assess cell and matrix phenotypes. Weekly time-lapsed micro-CT scans allow in situ monitoring of mineralization. (**D**) Postprocessing of micro-CT data with multiple thresholds and regional assessment of mineral densities and volumes. (**E**) Sample-specific micro-finite element analysis and mechanoregulation analysis allow investigation of the influence of the local mechanical environment on mineralization. Fig. created using BioRender.com.

## RESULTS

### Human bone-derived osteoblasts yield highly viable organotypic bone models

To fabricate patient-specific organotypic bone models, we isolated osteoblasts from surgical bone waste material from the femur (Fig. 2 A, B) based on adherence to tissue culture plastic, expanded and characterized the cells and created 3D bioprinted models. Osteoblasts from a 15-year-old metabolically healthy male and 3-year-old female OI patient homozygous for a pathogenic *FKBP10* c.890_897dup (p.Gly300Stop) variant showed comparable cell morphologies after isolation (Fig. 2E, I). Healthy osteoblasts cultured for 3 weeks in monolayer culture with osteogenic medium secreted a dense collagenous extracellular matrix (Fig. 2F, G), whereas osteoblasts from the OI donor showed reduced collagen deposition, a hallmark of *FKBP10*-related OI resulting from reduced collagen crosslinking (Fig. 2J, K) (*36*). We assessed cell viability after extrusion bioprinting (Fig. 2H, L), and found no significant decrease in viability in the OI-cell-laden scaffolds, with both healthy and OI groups exhibiting > 90% live cells (Fig. 2D).

**Fig. 2.**
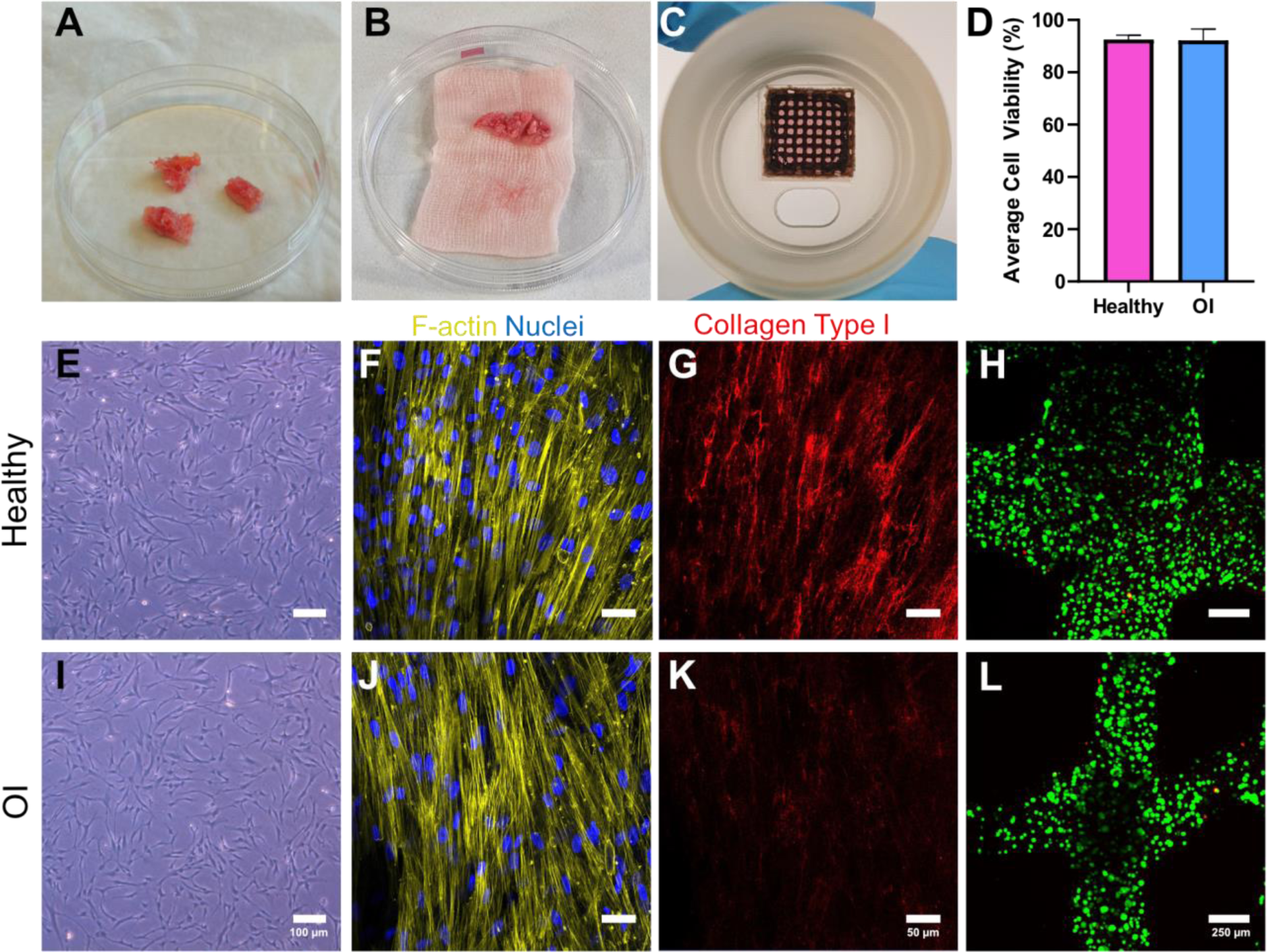
Biofabrication and characterization of patient-derived osteoblasts and organotypic bone models. (**A**) Bone material after orthopedic surgery of 15-year-old metabolically healthy male and (**B**) 3-year-old FKBP10-OI female (**C**) Brightfield image of extrusion 3D bioprinted organotypic bone model assembled in bioreactor (**D**) Average cell viability in 3D bioprinted healthy control and OI bone models. (**E**) Brightfield images of cell expansion taken on day 3 for (**E**) healthy and (**I**) OI osteoblasts. Scale bar = 100 µm. (**F**) Confocal imaging of actin filaments (yellow), nuclei (blue) of 2D patient cells after 3 weeks of culture in osteogenic medium for (**F**) healthy and (**J**) OI donors. Immunofluorescent staining overlay of the same region of interest showing collagen type I (red) for (**G**) healthy and (**K**) OI donors showing disease phenotype. Scale bar = 50 µm. (**H**) Representative fluorescent images of Calcein-AM/Ethidium homodimer-1-stained 3D bioprinted cell-laden scaffolds taken after bioprinting (day 1) for (**H**) healthy and (**I**) OI scaffolds. Live cells are green and dead cells are red. Scale bar = 250 µm.

### Time-lapsed micro-CT analysis shows increased TMD for OI organotypic bone models

To investigate tissue mineralization of the organotypic bone models, we used weekly micro-CT imaging (Fig. 3). Mineral volume (MV) was constantly higher in OI samples than in healthy samples (Fig. 3A, D), significantly higher from day 28 to 42 and becoming less pronounced at the end of culture. This is reflected in a significantly higher mineral formation rate for OI samples between day 14-21 and 21-28 (Fig. 3 B). Mineral formation rate was not significantly increased from day 28 onwards compared to healthy samples. Tissue mineral density (TMD) was significantly higher in OI than in healthy samples at day 42, 49, and 56 (Fig. 3C). Fig. 3D shows the mineralization process using time-lapsed images of a healthy and an OI sample. The 3D images underline that OI samples demonstrated more mineral volume than healthy samples over time, but that the end point MV was very similar. However, there were clear differences in how these samples mineralized. The healthy samples first mineralized in the top center of the scaffold and showed a more uniform mineralization from the center towards the sides and corners of the scaffold. The OI samples first mineralized at the scaffold sides in disconnected areas. In the center, MV formed around individual pores, demonstrating an inhomogeneous microarchitecture with sclerotic and void areas. TMD analysis confirmed this finding, with higher mineralized areas at the scaffold corners and around the pores than the healthy samples. The overall higher TMD of the OI sample was also clearly visible, with brighter greyscale intensities in the day 56 image compared to the healthy sample, especially at the scaffold surface. Overall, our analysis revealed not only faster (Fig. 3B) and higher (Fig. 3C) mineralization in OI samples than healthy controls, but also differences in the mineralization process and pattern between the groups (Fig. 3D).

**Fig. 3:**
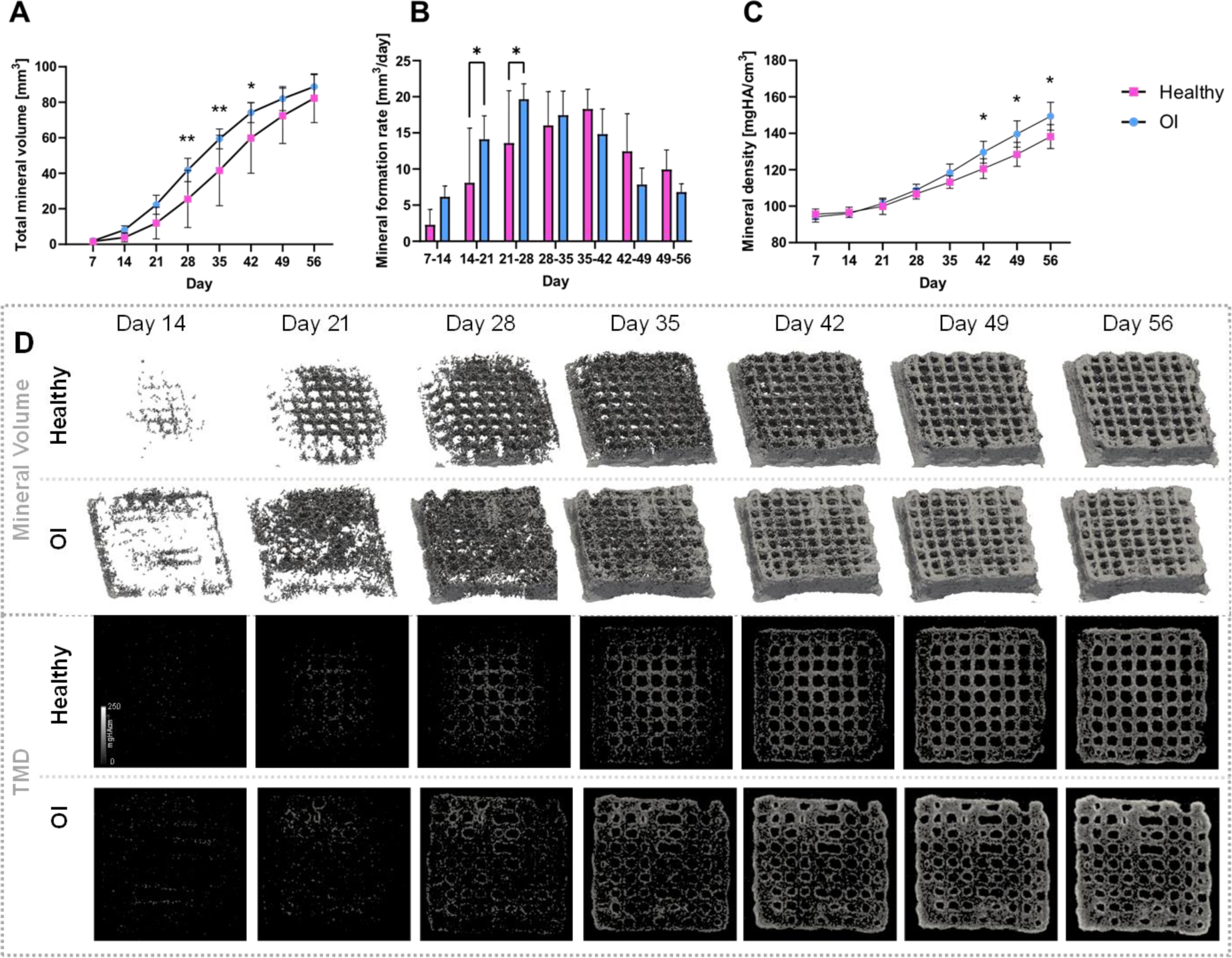
Time-lapsed analysis of mineralization reveals differences between healthy and OI organotypic models (n=6). **(A)** Mineral Volume, B: Mineral formation rate, **(C)** Tissue mineral density over time, **(D)** Qualitative time lapsed images of the mineralization process using 3D images of mineral volume and a 2D slice showing TMD of a healthy and an OI sample. Statistical significances among groups were identified by two-way ANOVA followed by Tukey’s post-hoc tests *p<0.05.

### Stiffness of OI organotypic bone organoids is not increased and can be predicted by micro-FE analysis

To investigate if the increased mineralization also led to increased stiffness in the OI samples, we determined compressive stiffness by uniaxial compression (Fig. 4C). However, OI organotypic bone model stiffness was not significantly increased compared to healthy controls, despite significantly higher TMD. To allow further insight into mechanical properties we conducted micro-FE analysis on the samples. By applying a top pad, we overcame the challenge of modelling realistic boundary conditions despite the uneven and variable top structure of the samples (Fig. 4A). Choosing a pad thickness for an embedded mineral volume of 2.2 mm^3^ (Fig. 4B) resulted in an R^2^=0.892 when correlating the resulting FE and experimental stiffness demonstrating FE analysis can predict scaffold stiffness for both healthy and OI samples (Fig. 4D). In Fig. 4E we plotted sample TMD against experimental and FE stiffness and compared the data with hMSC scaffolds from a previous study that showed a strong correlation between those two parameters. We observed that the healthy samples demonstrate a similar coupling of TMD and stiffness as the hMSC data. In contrast, OI samples do not match the original correlation, indicating a distinctive behavior compared to healthy samples and hMSC-laden scaffolds. Additionally, only one sample of OI scaffolds resulted in a higher stiffness than expected based on its TMD, while all other samples demonstrated a decreased stiffness from what the known correlation function would predict for their TMD. Additional results from micro-FE analysis from where no experimental data was available were in line with experimental results.

**Fig. 4:**
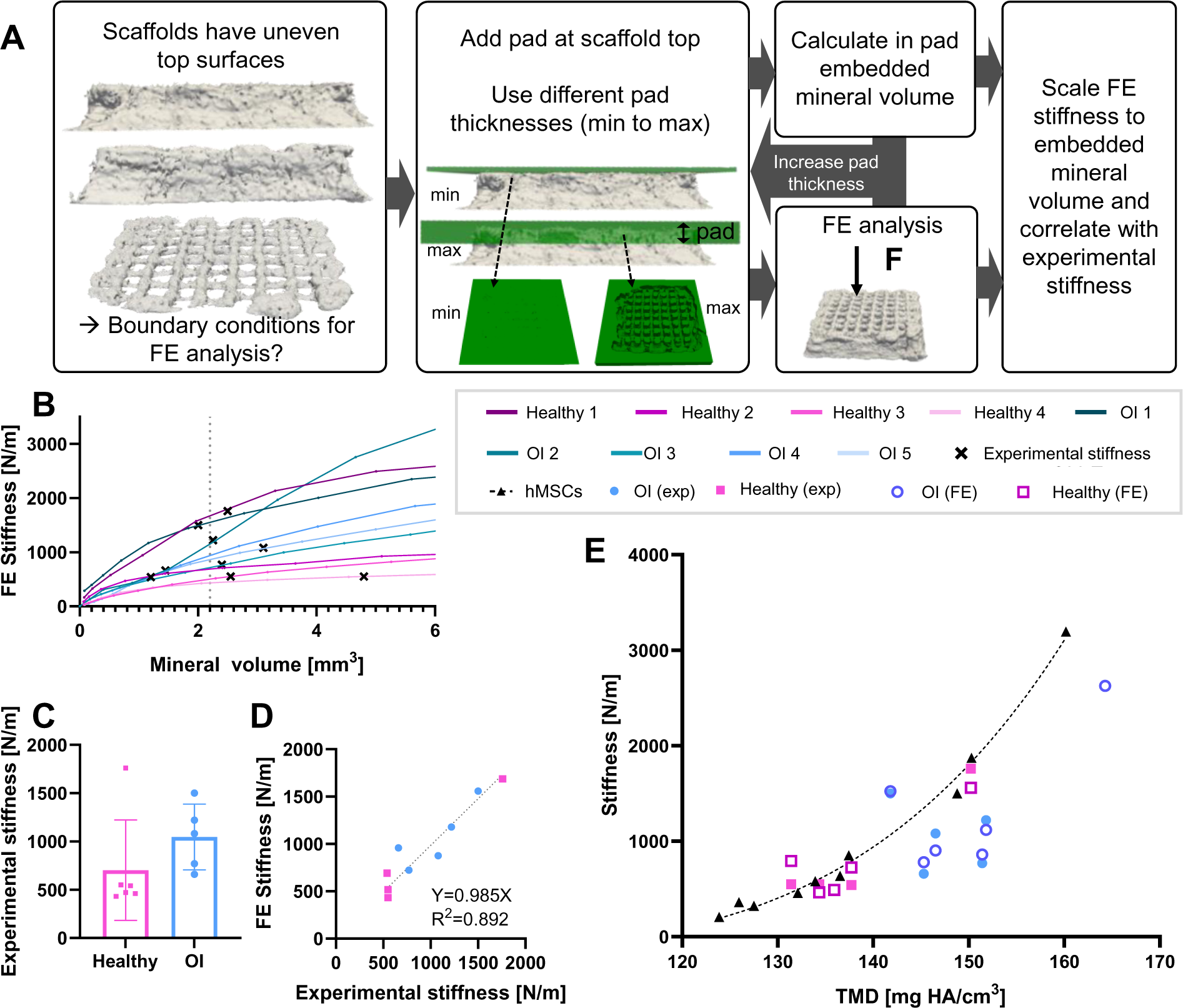
Experimental stiffness and stiffness calculated with micro-finite element (micro-FE) analysis of healthy and OI organotypic bone models. **(A)** Process for calculating stiffness with micro-FE analysis using a variable top pad to overcome difficult boundary conditions due to scaffolds uneven and variable top structure. **(B)** Comparing resulting FE stiffness with embedded mineral volume from the top pad to find a common value. **(C)** Experimental stiffness. **(D)** Correlation of experimental and FE stiffness using an embedded mineral volume of 2.2mm^3^. **(E)** Comparing experimental and FE stiffness with the underlying TMD and values from human mesenchymal stem cells (hMSC) scaffolds with a known TMD-stiffness correlation (39).

### Both healthy and OI organotypic bone models demonstrate mechanoregulation

To explore if dysregulated adaption to mechanical stimuli of the tissue formation process, called mechanoregulation, was responsible for the impaired stiffness of the OI scaffolds, we used a volumetric approach based on time-lapsed micro-CT images and micro-FE analysis to associate the mechanical environment with subsequent tissue formation (*40*) (Fig. 5). Our results demonstrated mechanoregulation in both groups (Fig. 5A), with no significant differences between healthy and OI samples in CCR or AUC (Fig. 5O,Q). Mean effective strain was significantly increased in tissue formation volumes compared with quiescent surface both in the healthy (45.72%, Fig. 5M) and the OI (47.09%, Fig. 5N) group, demonstrating tissue formation in regions of higher mechanical demand. Conditional probability curves of tissue formation also showed increasing probability of tissue formation with increasing effective strain on the scaffold surface for both groups (Fig. 5O). This can also be observed in the qualitative comparison of remodeling images and mechanical environment in Fig. 5A-L, especially in the close ups in Fig. 5I-L. The healthy samples show a slightly lower probability of formation at 0 effective strain, and slightly higher probability at 1 effective strain than OI, indicating mechanoregulation might be slightly increased compared to OI scaffolds. Correct classification rate (CCR) (Fig. 5P) and area under the curve for the receiver operating characteristic (AUC) for tissue formation (Fig. 5Q) also confirmed mechanoregulation for both groups, with slightly higher values for the healthy group, however not significant (CCR: p=0.42, AUC:p=0.21). Overall, our results demonstrate similar mechanoregulation in both healthy and OI organotypic bone models, and do not explain the impaired stiffness in OI samples.

**Fig. 5.**
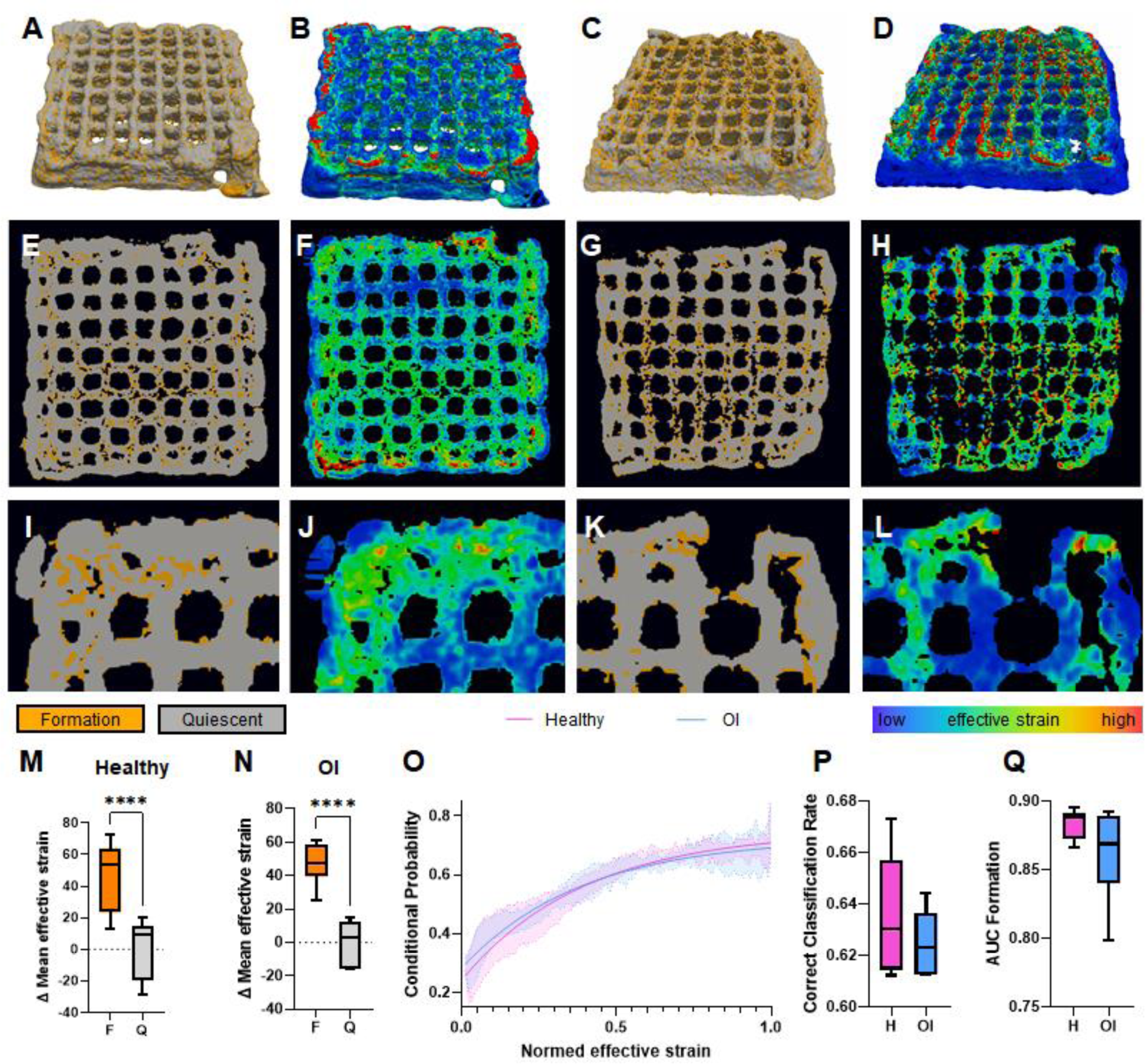
Mechanoregulation analysis in organotypic bone models and comparison between healthy and OI samples. **(A-L)** Qualitative comparison of mineralized tissue remodeling **(A,C,E,G,I,K)** and corresponding mechanical environment **(B,D,F,H,J,L)**, using 3D images of the whole scaffold **(A-D)**, a full 2D slice **(E-H)** and close ups **(I-L)** for one healthy example **(A,B,E,F,I,J)** and OI **(C,D,G,H,K,L)**. **(M,N)**: mean effective strain difference within formation volume compared to quiescent surface for healthy **(M)** and OI **(N)**. **(O)** Conditional probability curves of tissue formation dependent on the effective strain on the surface, **(P)**: Correct classification rate in healthy (H) and OI samples, **(Q)**: Area under the curve for the receiver operating characteristic for tissue formation. Significances calculated with paired (M,N) and unpaired (P,Q) students t-test, ****p<0.0001

### Regional analysis reveals differences in mineralization architecture for OI

To investigate the differences observed in the mineralization pattern between the OI and healthy samples, we divided the micro-CT images into 27 subsets by equally dividing the volume along each axis into 3 parts (Fig. 6E). The scaffold was divided into a top, middle, and bottom section. Due to the symmetry of the scaffolds, corner and side subsets were grouped and together with the center subset represent different areas within the scaffold plane. Since using the established threshold (84 mg HA/cm^3^) indicated marginal differences between groups, we implemented an additional high threshold (150 mg HA/cm^3^) for image segmentation to investigate highly mineralized mineral volume. Using this high threshold revealed differences between the samples (Fig. 6A, B). The healthy sample showed a continuous structure in the scaffold center and the bottom sides, but little highly mineralized tissue in the top corners (Fig. 6A). The OI sample had highly mineralized top corners, but the scaffold center consisted of both void and sclerotic volumes rather than a continuous structure (Fig. 4B). The regional analysis within the samples revealed no differences in TMD for healthy samples within the scaffold plane, except in the bottom (Fig. 6C). However, this was due to the center bottom subset showing very little mineralization in all samples (Fig. 6E). Other than this, the only regional differences in healthy samples were observed within scaffold height. In contrast to this, OI samples demonstrated more regional differences within the scaffold plane than within the scaffold height (Fig. 6D). Within a plane, corners showed the highest TMD (top: 155 mg HA/cm^3^, middle: 155 mg HA/cm^3^, bottom: 160 mg HA/cm^3^) and centers the lowest (top: 122 mg HA/cm^3^, middle: 132 mg HA/cm^3^, bottom: 103 mg HA/cm^3^), with only the difference between the middle side and center not being significant. However, within the scaffold height, only the bottom side showed increased TMD compared to the top side apart from the bottom center. This subset showed the lowest TMD among all subsets due to very little MV (Fig. 4E).

**Fig. 6:**
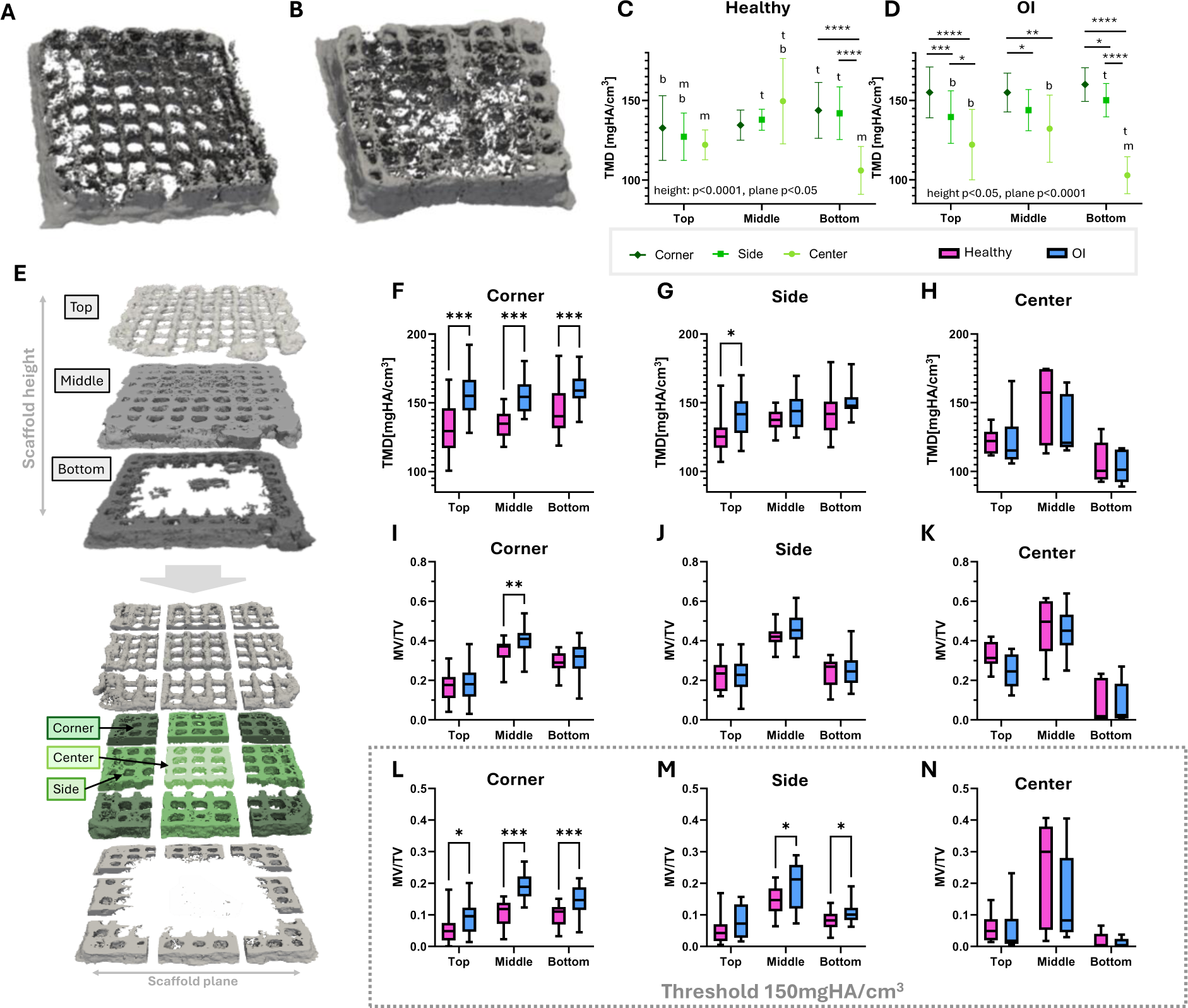
Regional mineral analysis of organotypic bone models using a low and a high threshold for image segmentation. Highly mineralized volume (using a threshold 150 mgHA/cm^3^ for image segmentation) of a healthy **(A)** and an OI **(B)** sample. **(C)** Regional analysis of TMD within healthy, and OI **(D)** samples. **(E)** Schematic explanation of the regional analysis using 27 subsets. Comparing regional TMD between healthy and OI samples for **(F)**: Corner, **(G)**: Side, **(H)**: Center subsets. **(I-K)** Comparing mineral volume per total volume (MV/TV) between healthy and OI samples for **(I)**: Corner, **(J)**: Side, **(K)**: Center subsets. **(L-N)** Comparing highly mineralized volume per total volume for **(L)**: Corner, **(M)**: Side, **(N)**: Center subsets. Statistical analysis for B, C with Two-way Anova with Tukey post-hoc test * p<0.05, ** p<0.01, *** p<0.001 for comparisons within the plane; t,b,m p<0.05 to top, middle, bottom for comparisons within scaffold height; F-N multiple unpaired t-tests with Holm-Bonferroni correction * p<0.05, ** p<0.01, *** p<0.001

Next, we compared similar regions between the two groups (Fig. 6F-N). Our analysis revealed that OI samples demonstrated consistently higher TMD in the scaffold corners across the whole scaffold height than the healthy control (Fig. 4F). While OI also showed higher TMD in the side subsets, the difference was only significant in the top subset (Fig. 6G). However, when comparing TMD in the center subsets, there were no significant differences between samples, and even slightly higher results for the healthy samples than for the OI, posing a contrast to the global results (Fig. 6H).

Mineral volume per total volume (MV/TV) was similar for both groups for sides and corners, except for the middle corner subset, where OI samples had significantly more mineral volume than healthy samples (Fig. 6I, J). In the center subsets, healthy samples showed slightly more mineral volume than OI samples, although not significant (Fig. 6K). However, applying a higher threshold revealed differences in volume of highly mineralized tissue (Fig. 6L-N). In line with previous results, we saw an increase in highly mineralized MV/TV in the corner subsets across the whole scaffold height for the OI samples compared to the healthy controls.

### Histological staining reveals phenotypic differences between OI and healthy controls

Next, we characterized the extracellular matrix of organotypic bone models to investigate the local mineralization and cellular environment after 8 weeks of uniaxial compression loading. Endpoint histological staining of consecutive sections of constructs revealed marked heterogeneity in the OI models. This heterogeneity, evident in the greyscale micro-CT slices of OI samples (Fig. 7E) corresponded to dense Alizarin Red S staining lining macroscale scaffold pores (Fig. 7N) and at scaffold corners (Fig. 7M). Additionally, OI samples exhibited increased microporosity within the printed struts (Fig. 7F-H, N-P). In contrast, healthy samples displayed more homogenous struts, particularly in central scaffold regions (Fig. 7J) with microporosity observed sparingly and localized towards the outer edges of the scaffold (Fig. 7K). This observation was supported by the mineralization pattern seen in the greyscale micro-CT slice, where the struts in healthy samples were found to be uniformly mineralized throughout the scaffold, with higher density (whiter) lining macroscale pores at the scaffold periphery (Fig. 7A), supporting the quantitative findings described in Figure 3 and 6.

**Fig. 7.**
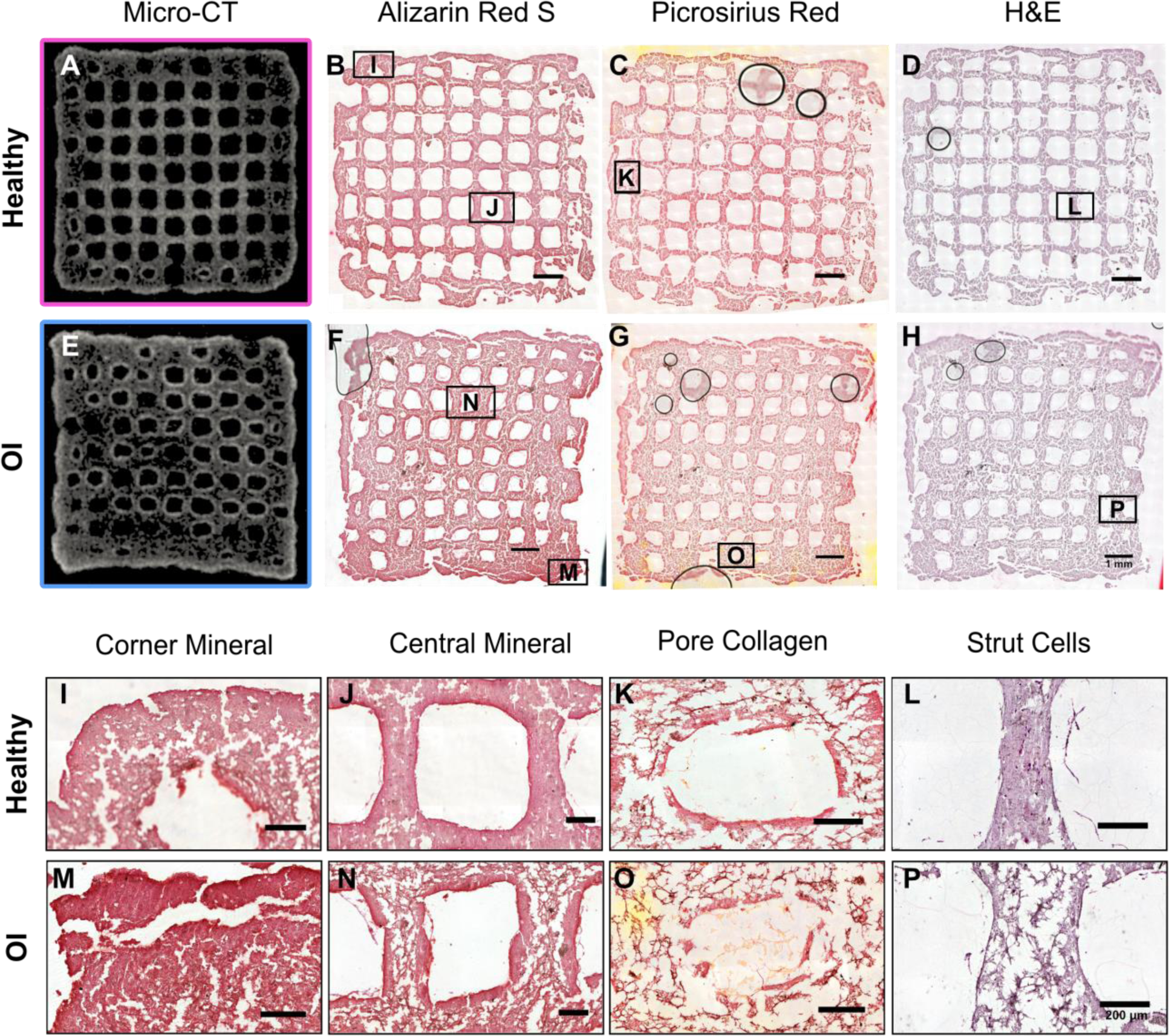
Characterization of Extracellular Matrix in Organotypic Bone Models. (**A**, **E**) Endpoint greyscale micro-CT slices reveal heterogeneity and voids in OI compared to homogenous mineral distributions healthy organotypic bone models. (**B**-**D**, **F**-**H**) histological staining of healthy (**A**-**D**) and OI (**E**-**H**) organotypic bone models indicates increased porosity in OI. Brightfield imaging of Alizarin red S (**F**) shows increased staining in corners (**M**) and lining pores of OI samples (**N**), Picrosirius red staining shows collagen (red-orange) in macroscale pores and surrounding embedded cells in both groups (**C**, **G**), Hematoxylin & eosin (H&E) staining shows cells spreading within porous printed struts and migrated cells in macroscale pores in both groups (**D**, **H**). Black circles are image artefacts caused by air bubbles. Scale bar = 1 mm. (**I**-**P**) Magnified regions of interest in healthy (**I**-**L**) and OI (**M**-**P**) organotypic bone models. Alizarin Red S staining of local mineralization in scaffold corners (**I**, **M**) and pores (**J**, **N**), Picrosirius red staining of collagen in pores (**K**, **O**) and H&E (**L**, **P**) staining of cell distributions in printed scaffold struts. Scale bar = 200 µm.

Hematoxylin eosin (H&E) staining revealed that cells were able to migrate out of the mineralized struts into macroscale construct pores in both healthy and OI groups (Fig. 7L, P). Particularly on the outer edges of the scaffold, where cell migration into the macroscale pores was more apparent, cell-secreted collagen matrices could be observed (Fig. 7K, O). Within scaffold struts in both groups, limited amounts of primarily peri-cellular collagen (stained orange around the embedded cells) was observed, indicating the potential of the developed organotypic model (Fig. 7K, O). Overall, histological staining revealed local differences in the porosity and mineralization of OI and healthy samples, supporting quantitative findings from regional micro-CT analysis (Fig. 6).

## DISCUSSION

In this study we developed a 3D-bioprinted patient-specific organotypic bone model mimicking mineralization dysregulation in *FKBP10*-related OI. To model healthy and pathogenic conditions, we created *in vitro* organotypic bone models with corresponding *in silico* micro-FE-models. Time-lapsed micro-CT imaging during culture revealed hypermineralization in the OI model compared to healthy controls, recapitulating aspects of the clinical disease phenotype. Despite higher mineralization, stiffness of OI samples was not significantly increased. We were able to show similar levels of mechanoregulation for both healthy and OI organotypic bone models, suggesting that impaired OI scaffold stiffness was not due to dysregulated mechanoregulation. A detailed regional analysis of TMD and highly mineralized tissue volume revealed increased heterogeneity of mineralization in OI scaffolds compared to healthy controls. Histological analysis confirmed microarchitectural inhomogeneities and indicated microporosity in scaffold struts and macroscale pores of OI samples, contrasting with the homogenous mineralization observed in healthy controls.

We found accelerated and overall increased mineralization in our OI models, which is consistent with findings in literature (*10, 13, 41*). While the corners of the OI scaffolds showed significantly higher mineralization than healthy controls, mineralization in the scaffold centers was not significantly different from healthy samples regarding TMD and mineral volume. This is in agreement with literature, where large variations in TMD have been observed in OI bone and cortical density can be both higher and lower compared to healthy controls in different sites (*21–23*). On a qualitative level, healthy samples showed a more homogeneous and coherent microarchitecture in the center compared to the OI samples, where we observed an inhomogeneous microarchitecture with both void and sclerotic parts. From a mechanical point of view, the microarchitecture in the center is important for an even load distribution and could be compared with trabecular bone architecture, which is known to be impaired in OI (*42, 43*). Bevers *et al.* recently reported similar observations of microarchitectural heterogeneity in trabecular bone in adults with OI using HR-pQCT (*24*), while Whittier *et al.* quantified comparable void spaces within osteoporotic trabecular bone which led to poor bone quality reflected in decreased bone strength (*25*). To analyze the influence of the observed dysregulation of mineralization on mechanics, we created FE models based on sample specific geometries with Youngs modulus derived from scaffold mineral density. Outputs from FE simulations reproduced the stiffness measured experimentally, using a correlation of TMD and Young’s modulus found in healthy tissue (*39*). This suggests regional differences in mineralization and heterogeneity are largely responsible for the impaired stiffness in OI, while material properties on a tissue scale are either comparable to healthy properties or their influence is minor. These results shed light on a long standing question of how much of impaired OI mechanics is caused by the mineral and how much by the organic component (*44*). Varga *et al.* also concluded that morphometrical parameters had a strong influence on mechanical parameters when using FE in an OI mouse model femur (Brtl/+) (*45*). Their prediction accuracy decreased slightly when using heterogenous material properties. Then again, homogeneous material properties are often used for *in vivo* work, while *in vitro* samples operate with a different range of Young’s modulus due to the comparatively low density and stiffness.

While physical therapy is recommended in OI disease management (*1*), mechanoregulation has not been specifically investigated yet. Whole-body vibration treatment was reported to increase trabecular bone volume and improve mechanical properties in a mouse model of severe OI (oim) (*46*). Similarly, Jeong and colleagues found that treadmill exercises increased femoral stiffness in +/G610C mice, modelling mild/moderate forms of OI (*47*). Additionally, using a myostatin inhibitor promoting muscle mass and thus mechanical loading in OI mouse models also led to increased bone morphometric parameters and partially to increased strength, indicating OI bone adapts to mechanical stimuli (*48*). This is in line with our finding of mechanoregulation in organotypic bone models of *FKBP10*-related OI. Conversely, a computational study investigating fluid flow in the lacuna-canalicular network found decreased streaming potentials and current in OI bone compared to healthy controls, suggesting that while OI bone remains susceptible to mechanical loading, its response may be diminished due to alterations in bone microstructure (*49*). Our findings emphasize the potential importance of physical therapy in OI disease management. Notably, studies on osteoporosis suggest that the type of exercise modality significantly influences the effectiveness of physical therapy as an anabolic treatment and in reducing fracture risk (*50*). Hoyer-Kuhn and colleagues (*51*) demonstrated that a comprehensive rehabilitation program, incorporating whole body vibration, resistance training, and treadmill running, led to improved mobility and bone mineral density compared to whole body vibration alone (*52*).

The organotypic bone models showed limited collagen deposition, a phenotype previously reported in primary osteoblast constructs (*53*), posing a limitation of this study. *In vitro* models using encapsulated mesenchymal stem cells or iPSCs as starting cells have been shown to produce collagen (*37, 39*). However, patient-specific osteoblasts offer the advantage of shorter protocols avoiding the establishment of cell lines, risk of off-target cell types or epigenetic changes from cellular reprogramming, fitting into the existing clinical pipeline for OI care (*37, 38, 53–55*). Moreover, studies have suggested hypermineralization and other mineral alterations in OI occur independent of the collagen mutation (*11*) and our goal was to focus on those alterations of mineralization in a controlled environment. Due to collagen’s role in bone elasticity, this model may not represent the toughness and brittleness phenotypes of OI. However, Varga *et al.* concluded morphology as a much more dominant determinant of bone strength than intrinsic bone properties using FE analysis, indicating that including collagen would have limited impact on global mechanics (*45*). Secondly, the donors used in this study were not age or gender matched, but Gamsjaeger *et al.* reported only minor age and sex related differences within healthy children (*56*). Additionally, Rauch *et al.* found that within one donor woven bone, recently remodeled tissue, and lamellar bone properties were more different than similar regions between donors of different age and sex (*21*). Lastly, the studies used a relatively small sample size and detected high intragroup variability. However, this reflects clinical variability, thus, our model’s ability to mimic this heterogeneity using cells from a single donor can also be considered a strength.

We have successfully developed a model of dysregulated mineralization in an *FKBP10*-related OI organotypic bone model possibly enabling screenings of treatments as preselection assessment for animal studies or clinical trials, accelerating the search for targeted OI therapies. The organotypic bone model could help investigate the mechanism of action of novel therapies minimizing the ethical concerns associated with animal models, providing a tunable, humanized environment yielding personalized constructs within weeks. Additionally, creating a second model incorporating collagen would allow us to explore further mechanical aspects such as brittleness and toughness, shedding light on how treatment can influence these properties. Finally, the goal is to broaden the donor cohort by establishing a biobank to include more patients incorporating different OI mutations. Together, these results suggest this organotypic bone model may be a promising tool to study pathological mineralization mechanisms and, in the future, assess personalized disease progression and treatments in OI.

## MATERIALS AND METHODS

### Study Design

In this study, we describe a novel *in vitro* approach for studying osteogenesis imperfecta. Primary osteoblasts were isolated from surgical waste materials of pediatric patients. By extrusion 3D bioprinting organotypic bone models and monitoring mineral maturation through non-destructive *in situ* time-lapsed micro-CT imaging, we were able to evaluate bone development under physiological loading conditions. By applying multi-density thresholding and dividing the micro-CT scans into subsets, mineral volume and mineral formation rate in diseased states can be investigated. By combining computational and experimental techniques we demonstrate *FKBP10*-OI organotypic bone models resemble some aspects of the disease *in vitro*.

### Ethic Statement

The studies involving human participants were reviewed and approved by Swiss Ethics (KEK-ZH-Nr. 2019-00811 to CG and MaR). Written informed consent to participate in this study was provided by the participants’ legal guardian/next of kin.

### Isolation of primary osteoblasts

Bone explants were collected from an intermedullary rodding surgery of a 3-year-old female donor (*FKBP10* c.890_897dup (p.Gly300Ter)) and a femur osteotomy of a healthy 15-year-old male donor with limb malalignment as waste material under the study protocol approved by Swiss Ethics (KEK-ZH-Nr. 2019-00811 to CG and MaR). Upon collection, bone explants were immersed in Dulbecco’s Modified Eagle Medium (DMEM, Gibco), washed in Phosphate Buffered Saline (PBS, Gibco), cut into approximately 10 to 20 mm long pieces, and then vortexed in fresh PBS to remove blood contaminants. The bone explants were transferred to tissue culture dishes and cultured at 37 °C and 5 % CO2 in DMEM supplemented with 10% fetal bovine serum (FBS, Gibco), and antibiotic-antimycotic (Gibco, containing 100 U/ml penicillin, 100 mg/ml streptomycin, and 0.25 mg/ml Amphotericin B). The explants were left undisturbed for 7 days, and the culture medium was changed every 3 to 4 days thereafter. Cells that migrated out of explants and attached to the culture dishes were dislodged by trypsinization and expanded in T75 culture flasks until they reached 90 % confluency, after which they were cryopreserved in FBS and 10 % dimethyl sulfoxide (DMSO).

### Characterization of primary osteoblasts

Osteoblasts were characterized under standard 2D cell culture conditions. DNA of the mutated gene region (exon 5 of the *FKBP10* gene) was amplified (5’ to 3’ sequence forward primer GGG CTA GTG TCT TGC ATG GT and reverse primer CAG CTT CGT CCA CGT TTC AC), followed by gel electrophoresis and capillary Sanger sequencing.

Hallmarks of collagen disorder were assessed by IHC staining of collagen ECM. Primary osteoblasts from control and *FKBP10*-OI patients were cultured on chamber slides (µ-Slide 2 Well-ibiTreat, ibidi). To support cell adhesion, cells were cultured for three days in control medium before supplementing with osteogenic medium for 3 weeks to promote ECM deposition. Samples were washed twice in PBS and fixed in 4% paraformaldehyde (PFA), washed in PBS, and subsequently stained with anti-collagen I antibody (ab34710, Abcam). The samples were then incubated with secondary donkey anti-rabbit AF647 antibody (1:1000, ab150075, Abcam). Samples were permeabilized and blocked in 1% BSA and 0.1% Triton-x-100 and subsequently stained with Phalloidin-TRITC (1:1000, P1951, Sigma-Aldrich) and Hoechst 33342 (1:200, 14533, Sigma-Aldrich). Samples were imaged using a confocal microscope (Zeiss LSM 880 Airyscan, Germany) and postprocessed in Fiji.

#### Organotypic bone model biofabrication

##### Preparation of bioinks

Primary osteoblasts (passage 5) were harvested by incubation with 0.25 % Trypsin-EDTA and resuspended in the control medium (DMEM, 10 % FBS, 1 % Anti-Anti). These cell suspensions were maintained at 4 °C and, as needed, underwent centrifugation, followed by resuspension in 60 μL of control medium. The cell suspensions were combined with 1 ml of hydrogel mixture comprising 4.1% (w/v) gelatin, 0.8% (w/v) alginate, and 0.1% (w/v) graphene oxide, following established procedures (*57*).

##### Extrusion 3D bioprinting

Bioinks containing 10×10^6^ cells/mL were loaded into 3-ml polyethylene cartridges equipped with 27-gauge tapered tips (Nordson EFD, Vilters, Switzerland). The 10 mm × 10 mm × 2.4 mm scaffolds were printed using a 3DDiscovery bioprinter (RegenHU; Villaz-St-Pierre, Switzerland) with a pneumatic dispenser onto double-sided tape (3M, Scotch, USA) affixed on the bioreactor platform, as previously described (*39*). Scaffolds were submerged in 2 % (w/v) calcium chloride in control medium for 10 minutes to crosslink the hydrogel, then washed twice in control medium. Scaffolds were transferred to 6-well plates with fresh control medium and incubated at 37°C with 5 % CO_2_.

##### Dynamic culture

Scaffolds were assembled into custom-made polyetherimide compression bioreactors. Each bioreactor was filled with 5 ml osteogenic medium (DMEM, 10 % FBS, 1% Anti Anti, 50 µg/ml ascorbic acid, 100 nM dexamethasone, 10 mM β-glycerophosphate) with media changes performed three times per week. Scaffolds underwent cyclic loading in a mechanical stimulation unit (MSU) controlled via a custom LabView program (National Instruments, Austin, Texas). The loading protocol consisted of uniaxial compression loading with a preload (0.07 N) to ensure piston-sample contact, followed by sinusoidal compression (1% strain, 5 Hz, 5 minutes). Organotypic bone models were mechanically loaded 5 times per week enabling maturation over 8 weeks.

#### Characterization of cells and extracellular matrix (3D)

##### Cell viability

Cell viability in extrusion 3D bioprinted scaffolds was evaluated with LIVE/DEAD® Viability/Cytotoxicity assay 24hrs post bioprinting. In brief, scaffolds were incubated with 2 μM Calcein AM and 4 μM ethidium homodimer for 40 min 37 °C and 5% CO_2_. Following this, scaffolds were washed twice with pre-warmed PBS and transferred to 8-well chamber slides (Ibidi GmbH, Germany) for imaging using a confocal microscope (Visitron Spinning Disc, Nikon Eclipse T1). Cell viability was calculated using ImageJ (National Institutes of Health, USA) as the ratio of living cells to the overall cell count.

##### Mechanical testing

Scaffold mechanics were evaluated using the in-house MSU as described previously (*39, 58*). Unconfined uniaxial compression tests were conducted under displacement control, employing a preload of 0.07 N, and a displacement rate of 4 µm/s until the scaffold yielded. Subsequently, stiffness was calculated from the linear region of the resultant force–displacement curve.

##### Sample preparation, histological staining and imaging

On day 56 of the study, organotypic bone models were carefully removed from their bioreactors and fixed in 4% PFA for 2 h and subsequently immersed in 10% sucrose solution for 2 h, followed by overnight incubation in 30% sucrose solution. Scaffolds were embedded in optimal cutting temperature compound (OCT, VWR) in a methanol bath on dry ice and subsequently stored at - 80° C. Samples were sectioned (10-30 µm thickness) using Kawamoto’s cryofilm type 2C (SECTION-LAB Co. Ltd., Japan) using a cryotome (CryoStar NX70, Thermo Scientific) (*59*). Prior to staining, sections were affixed to microscope slides (SuperFrost™ Microscope Slides, ThermoScientific) using a solution of 1% (w/v) chitosan in 1% (v/v) acetic acid.

To facilitate the visualization of different components, a series of staining techniques were employed. Hematoxylin (Mayer’s, Sigma-Aldrich) and eosin Y disodium salt (Sigma-Aldrich) (H&E) staining was performed to visualize cell nuclei, cytoplasm, and extracellular matrix.

Alizarin Red S staining (2mg/ml in acetone pH 4.3) (A5533, Sigma-Aldrich) was used to stain the mineralized extracellular matrix. Picrosirius red staining (365548, P6744, Sigma-Aldrich) enabled visualization of collagen. The histological sections were imaged with an automated slide scanner (Panoramic 250 Flash II, 3Dhistech, Hungary) at 20x magnification.

##### Immunohistochemistry staining and confocal imaging

Cryosections of organotypic bone models were washed thrice with NaCl/HEPES buffer (135 mM NaCl, 20 mM HEPES, 10 mM NaOH, pH 7.47), blocked and permeabilized with 5% BSA in NaCl/HEPES-0.1% triton-x100 for 1 h. Samples were stained overnight with anti-collagen I antibody (1:200, ab34710, Abcam) in 1% BSA in NaCl/HEPES. Samples were washed and incubated with secondary donkey anti-rabbit AF647 antibody (1:1000, ab150075, Abcam) and subsequently stained with Phalloidin-TRITC (1:1000, P1951, Sigma-Aldrich) and Hoechst 33342 (1:200, 14533, Sigma-Aldrich). Samples were imaged using a confocal microscope (Zeiss LSM 880 Airyscan, Germany) and postprocessed in Fiji.

##### Real-time quantitative polymerase chain reaction

Total RNA was extracted from organotypic bone models cultured in osteogenic medium for 2, 3, and 4 weeks under static conditions, according to the manufacturer’s protocol (RNeasy Mini kit, Qiagen). The isolated RNA was reverse transcribed to cDNA following the manufacturer’s protocol (PrimeScript™ RT Master Mix, TaKaRa). The mRNA levels of RUNX2 (Hs00231692_m1), COL1A1 (Hs00164004_m1), COL1A2 (Hs01028956_m1), BGLAP (Hs01587814_g1), PDPN (Hs00366766_m1), DMP1 (Hs01009391_g1) and SOST (Hs00228830_m1) TaqMan probes (TaqMan Gene Expression Assays, Applied Biosystems) were detected by real-time quantitative polymerase chain reaction (CFX96™ Real-Time System, Bio-Rad). Relative gene expression was normalized to housekeeping gene glyceraldehyde-3-phosphate dehydrogenase (GAPDH) (Hs02758991_g1).

#### Computational analysis

##### Time-lapsed micro-computed tomography

Bioreactors were scanned every 7 days in a micro-computed tomography (micro-CT) scanner (µCT45, SCANCO Medical AG, Brüttisellen, Switzerland) at a voxel resolution of 34.5 µm with an energy of 45 kVp, intensity of 177 µA, and an integration time of 600 ms. Micro-CT data was evaluated with VMS and Python (Python Software Foundation, Delaware, USA) (*57, 58*).

##### Image processing

Time-lapsed micro-CT images were registered to the day 7 image for each sample and gauss-filtered (sigma:1.2, support 1.0). Images were segmented using a threshold of 84 mgHA/cm^3^ (*39*), and TMD was calculated from segmented greyscale images. Binarized segmented images were used to calculate total mineral volume and mineral formation rate, by subtracting the mineral volume from the previous week for each sample. 3D images of mineral volume and 2D images of TMD were visualized in Paraview (Kitware, Clifton Park, New York).

##### Micro-Finite Element Analysis

Segmented greyscale images of day 49 and 56 were converted to Young’s Modulus using the conversion function published by Zhang *et al.* and voids within the scaffolds were assigned a Young’s modulus of 3kPa to account for not mineralized connecting structures (*39, 40*). Due to the uneven and soft top of the scaffolds, which varied between samples, we implemented a top pad with a Young’s Modulus of 5GPa with multiple thicknesses adapted to the scaffold geometry (Fig. 4A), enabling more realistic boundary conditions and force flow (*40, 60*). Padding thickness was adjusted so that it ranged from minimal contact with the scaffold to a contact area of 70% of the maximum cross sectional area of each sample, increasing one voxel in thickness at a time. A 1% uniaxial compression was applied to the top nodes (*60*). Micro-FE analysis was performed using the linear solver ParOsol (*61*) with 1 node and 12 cores for each padding thickness and sample averaging in 40 million elements, and converging in 2 minutes, returning the effective strain (*62*) and resulting force. Stiffness was calculated based on the resulting force and sample height. To enable a comparison between samples despite varying geometry and to choose the most appropriate pad thickness amongst the investigated, we scaled the resulting stiffness (day 56) to the embedded mineral volume by the respective pad thickness (Fig. 4B). We then correlated the resulting stiffnesses to experimental values between 0 and 6mm^3^ and a mineral volume of 2.2 mm^3^ was chosen as a common denominator with the best correlation between experimental and micro-FE stiffness (0.89). The final micro-FE stiffness and subsequent analysis was based on the results using a top pad with an approximately embedded mineral volume of 2.2mm^3^. Day 56 images were used for the stiffness analysis, while day 49 images were used for the mechanoregulation analysis.

##### Mechanoregulation Analysis

Mechanoregulation analysis was performed based on the resulting effective strain from micro-FE analysis of the micro-CT images of day 49 and a remodeling image labeling formed and quiescent tissue between day 49 and 56. For this, registered, segmented and binarized images of day 49 and 56 were superimposed, and tissue present in both images was labelled quiescent and tissue only present on day 56, but not on day 49 was labelled formation. To associate effective strain from micro-FE analysis with tissue formation, we applied a previously developed method, which uses a volumetric projection of surface effective strain into the formation volumes (*40*). Mean effective strain of formation volumes was calculated and compared to mean effective strain on the quiescent surface. Additionally, conditional probability curves for tissue formation were calculated as described elsewhere (*40, 63*). AUC for tissue formation was computed based on formation associated effective strain and effective strain on the scaffold surface, resulting in a value between 0.5 and 1, indicating the level of mechanoregulation present in tissue formation (*64, 65*). Similarly, CCR was calculated, indicating the level of overall mechanoregulation, also a value between 0.5 and 1 (*66, 67*). For both AUC and CCR, a value of 0.5 indicates no dependency of formation on the mechanical environment, while a value of 1 indicates formation is completely controlled by the mechanical environment.

##### Regional analysis of mineralization

For the regional analysis, segmented greyscale micro-CT images of day 56 were divided into 27 subsets by equally splitting the image into thirds along each axis (3×3×3). To ensure comparability between samples, micro-CT images were visually checked for alignment of scaffold with image axes and manually corrected if necessary. Additionally, image shape was adjusted so that there were no empty voxel layers on the image outsides. Within each subset, TMD and MV/TV was calculated. Additionally, we calculated MV/TV for highly mineralized tissue using a threshold of 150mgHA/cm^3^. For terminology, we defined scaffold height as the z-axis and scaffold plane as the x-y plane. Due to the symmetry of the scaffold within the plane, we grouped corner and side subsets within a scaffold plane and sample (Fig. 6E), leading to 9 subset categories: top corner, top side, top center, middle corner, middle side, middle center, bottom corner, bottom side, and bottom center. We first compared subset TMD within the healthy and the OI samples to analyze regional mineralization heterogeneity. Second, we investigated differences between healthy and OI samples by comparing TMD, MV/TV and highly mineralized MV/TV among healthy and OI samples.

#### Statistical Analysis

Statistical analysis was performed using GraphPad Prism 10 software. P-values less than 0.05 were considered statistically significant. Data are represented as mean ± standard deviation. The comparison of cell viability was performed using an unpaired two-tailed Student’s t-test. The comparison of micro-CT data between groups at different timepoints was done using a two-way ANOVA followed by Tukey’s post-hoc test. For the mechanoregulation analysis, data were first tested for normality. Statistics between mean effective strain was calculated using paired Student’s t-test, for CCR and AUC values using unpaired students t-test. For the regional analysis, data were first tested for normality using Shapiro-Wilk-test. Differences within a group were analyzed using two-way ANOVA followed by Tukey post-hoc test. Differences between OI and healthy subsets were calculated with multiple unpaired Student’s t-tests and Bonferroni-Holm correction for multiple comparisons.

## List of Supplementary Materials

Fig. S1-3

## Acknowledgments

The authors thank the Scientific Center for Optical and Electron Microscopy (ScopeM) of ETH Zurich for providing microscopy instrumentation and support.

## Author contributions

Conceptualization: JG, ADL, MR, MR, CG, RM

Methodology: JG, ADL, PJL, BK, TN, MR

Investigation: JG, ADL, TM, BK, TN, PJL

Visualization: JG, ADL, TM, BK

Supervision: SJE, MJM, JLS, EH

Writing – original draft: JG, ADL

Writing – review & editing: JG, ADL, FS, RM

