## Supplementary Material for "3D-bioprinted patient-specific organotypic bone model mimicking mineralization dysregulation in *FKBP10*-related osteogenesis imperfecta"

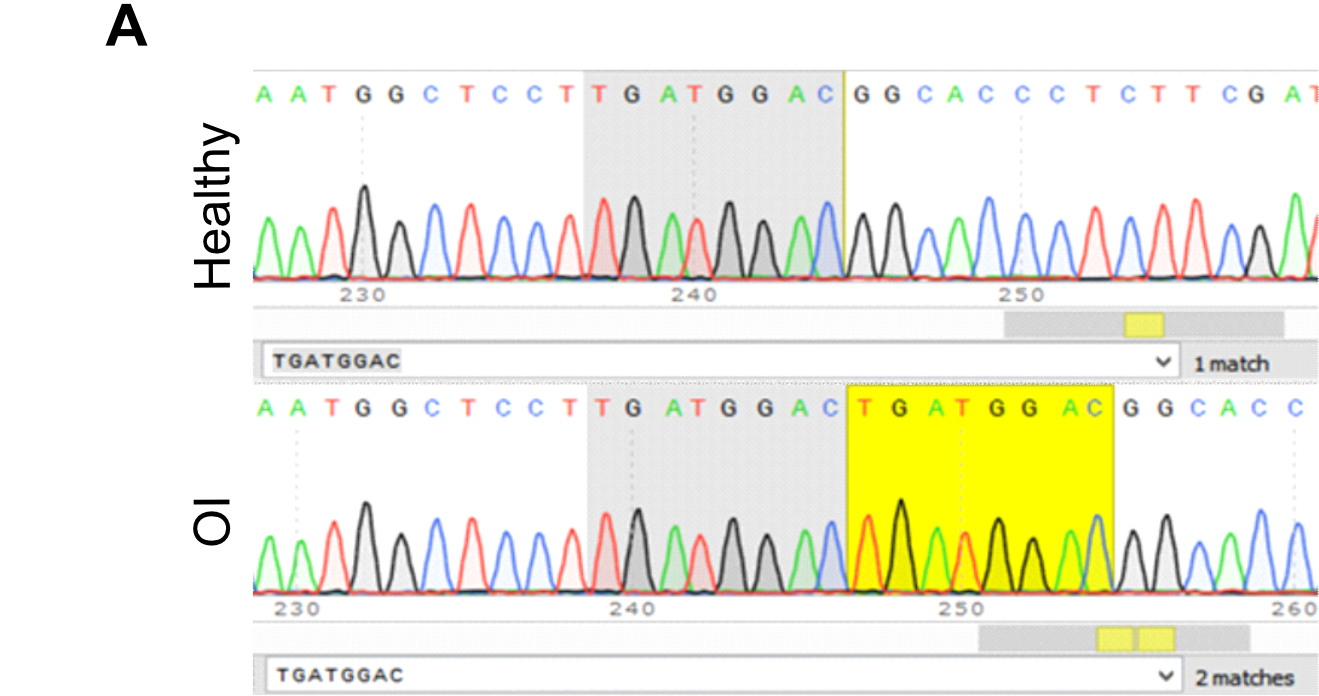

***Fig. S1. Confirmation of OI mutation***

*(****A****)* *Comparison of DNA sequencing chromatograms of healthy and OI donors. Sanger sequencing of exon 5 of the FKBP10 gene revealed the homozygous duplication of the 8 base pair TGATGGAC sequence in the OI cells, corresponding to the c.890_897dup (p.Gly300Ter) mutation.*

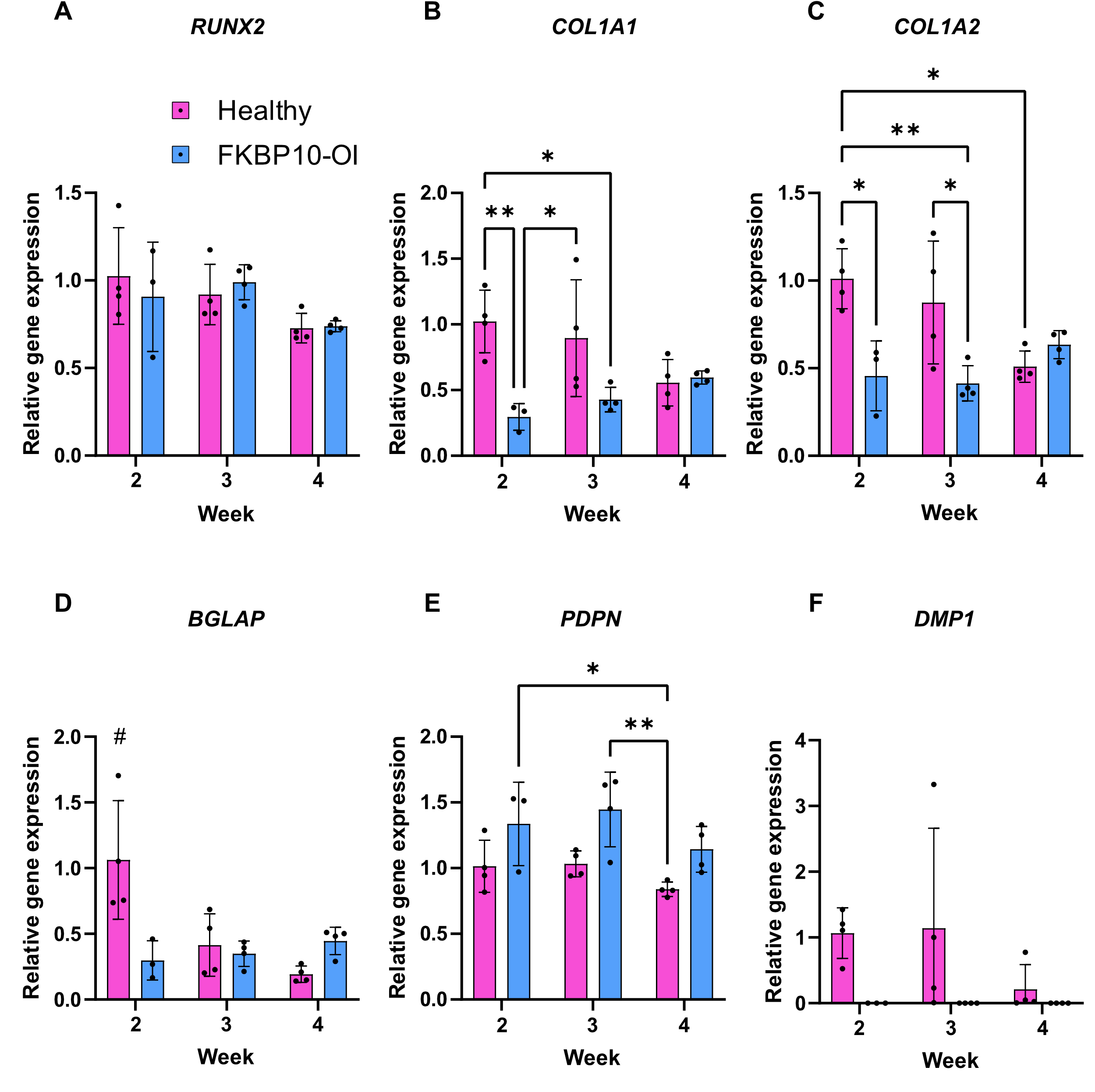

***Fig. S2. Relative osteogenic gene expression in static organotypic bone models.***

*(****A****) Quantitative RT-PCR analyis of Runt-related transcription factor 2 (RUNX2), (****B****) collagen type I alpha 1 chain (COL1A1), (****C****) collagen type I alpha 2 chain (COL1A2), (****D****) osteocalcin (BGLAP), (****E****) podoplanin (Pdpn), and (****F****) Dentin matrix acidic phosphoprotein (DMP1) for healthy and OI organotypic bone models after 2, 3, and 4 weeks of static culture in osteogenic medium normalized to healthy week 1 data. * P<0.05, # p<0.05 compared to all other groups.*

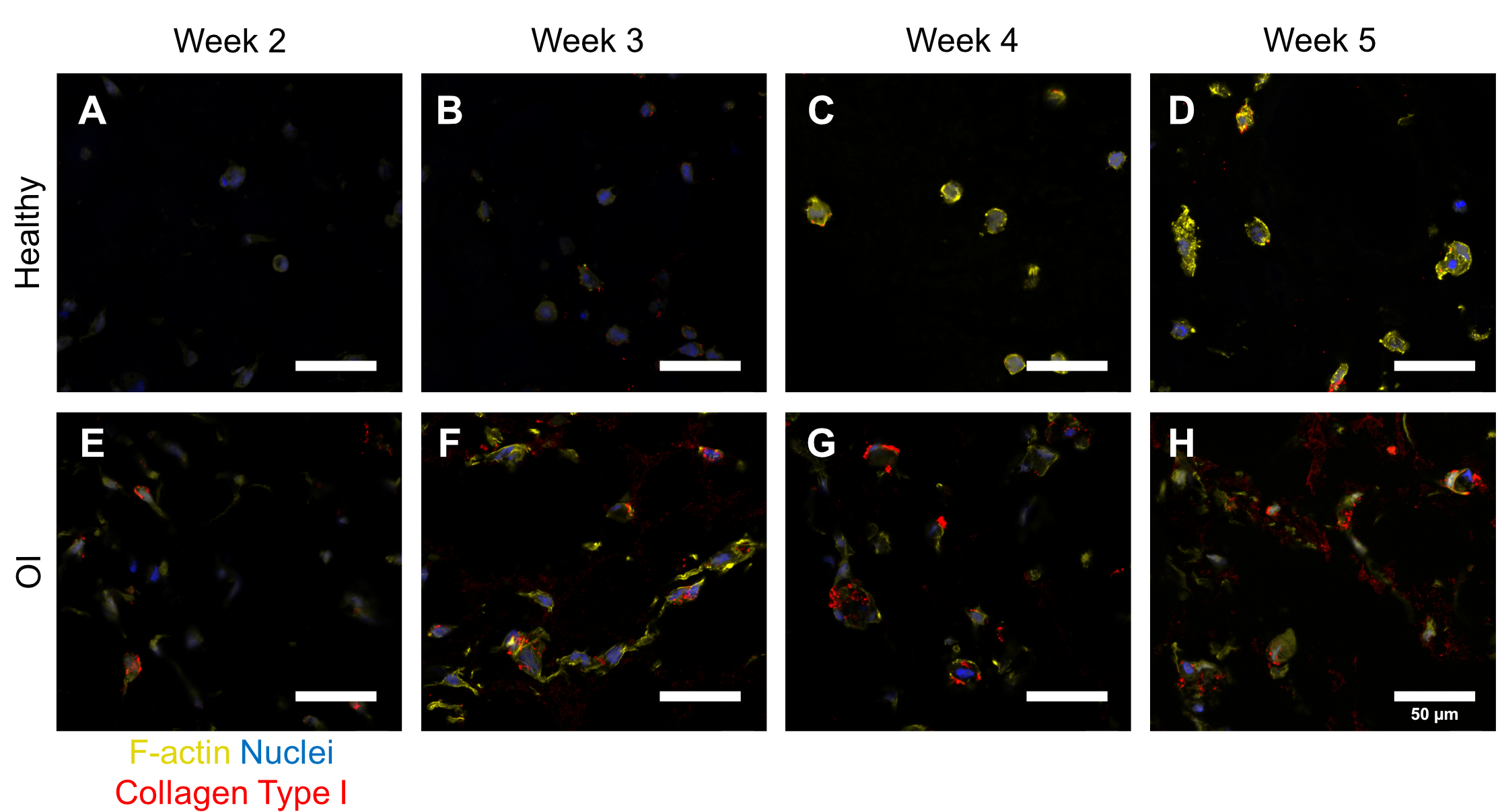

***Fig. S3. Relative osteogenic gene expression in static organotypic bone models.***

*(****A****) Immunofluorescent staining of actin filaments (yellow), nuclei (blue), collagen type I (red) in cryosections of organotypic bone models after 2 to 5 weeks of static culture in osteogenic medium for (****A****-****D****) healthy and (****E****-****H****) OI groups. Scale bar = 50 µm.*
